# Surgery and Radioactive Iodine Therapeutic Strategy for Patients Greater than 60 Years of Age with Differentiated Thyroid Cancer

**DOI:** 10.1101/2020.03.13.990341

**Authors:** Tao Tang, Wei Zhang, Jingtai Zhi, Xianhui Ruan, Linfei Hu, Xiaoyu Chen, Zhaohui Wang, Xiangqian Zheng, Ming Gao

## Abstract

**Purpose:** The purpose of the current study was to determine whether older patients with differentiated thyroid cancer (DTC) who received surgical treatment had a better cause-specific survival (CSS) than patients who were recommended surgery, but declined, and whether patients who underwent post-operative RAI-131 therapy had an impact on CSS based on TNM staging and number of lymph node metastases for all total or near-total thyroidectomy patients.

**Patients and Methods:** All DTC patients information were obtained from the SEER*Stat 8.3.6 program, and only patients ≥ 60 years or older were considered. The patients were divided into two groups (underwent surgery and surgery recommend, but not performed). Furthermore, patients were grouped as follows: T4; N1b; M1; T1-3N0-1a; specific number of lymph node metastases; and total or near-total thyroidectomy.

**Results:** The 120-month cause-specific survival (CSS) rate of females and males showed a gradual declining trend from 60-64 to ≥80 years of age in the group that underwent surgery. The CSS rate of females and males showed a marked downward and irregular trend with an increase in age in the recommended, but no surgery group. Univariate analysis indicated that the surgery group had a higher 120-month CSS in females in most stages and males, compared with the no surgery group. RAI-131 therapy was associated with an improved 80-month CSS in T4/N1b/M1 females (P<0.0183) and males (P<0.0011). There was no CSS difference in females or males between the T1-3N0 and T1-3N1a patients. There was no statistical difference between the two subgroups.

**Conclusions:** Surgical treatment should be recommended for elderly DTC patients because surgery can lead to a better CSS. High-risk patients achieve a higher benefit-to-risk ratio with RAI-131 therapy. To avoid the adverse effects associated with RAI-131 therapy, a multidisciplinary discussion should be arranged for intermediate- and low-risk patients.

## Introduction

Thyroid cancer is the most frequent endocrine malignancy and the incidence has nearly tripled in the past few decades in the US [1]. Among thyroid cancers, > 90% are differentiated thyroid cancer (DTC), 5% are poorly differentiated thyroid cancer (PDTC), 1% are anaplastic thyroid cancer, < 3% are medullary thyroid carcinoma [2]. The incidence of thyroid cancer is approximately 4-fold higher in women than men. The significant increase in the incidence of thyroid cancer is ongoing and resulted in thyroid cancer becoming the third most common cancer in women of all ages by 2019 [3].

Thyroid cancers have a global mortality rate of < 2% at 5 years. The 5-year mortality rate of thyroid cancers was < 2 % and remained stable from 1975-2009 (approximately 0.5 deaths per 100,000). With the gradual increase in the elderly population and the popularization of physical examinations, more and more elderly patients with DTC have been diagnosed.

The standard treatment of DTC is a radical operation, but some patients decline to undergo surgery and surgery is not recommended for other patients with severe diseases. In this study we determined whether older thyroid cancer patients who underwent surgical treatment had a better cause-specific survival (CSS) than patients who were recommended to undergo surgery, but declined, without any pre-existing assumption or bias.

Post-operative disease status should be taken into consideration when deciding whether to recommend additional treatment. In general, post-operative radioactive iodine-131 (RAI-131) adjuvant therapy is routinely recommended for DTC patients at high-risk based on the American Thyroid Association (ATA) guidelines. RAI-131 therapy has been used for patients for the following indications: remnant ablation; potential micro-metastases; loco-regional invasion; and pulmonary or bone metastatic disease. The purpose of therapy is to ensure a tumoricidal effect [4,5]. Treatment has been recommended more selectively in recent years as guidelines have evolved to reflect risks and utility in patient subsets. However, is treatment suitable for all patients with lymph nodes metastases and can treatment reduce the risk of death or recurrence?

We sought to determine the outcomes between patients who underwent surgery and patients in whom surgery was recommended, but declined. We also determined the outcomes between elderly patients with DTC who did and did not receive post-operative RAI-131 therapy. We investigated the comparative effectiveness of these patients by measuring CSS and the duration of survival in patients of different ages and stages based on the Surveillance Epidemiology and End Results (SEER) Medicare-linked database.

## PATIENTS AND METHODS

### Study Sample and Data Sources

All data regarding demographic characteristics and cancer incidence were obtained from the SEER*Stat 8.3.6 program of incident cases from 1975-2016 (November 2018 submission) [6].SEER is the primary source for cancer statistics in the United States, and represents 28% of the US population [7]. SEER is considered as representative of the US in terms of demographic composition, as well as cancer incidence and mortality rate.

### Patient Selection

We selected all patients who met the following criteria (Figure 1): site recode ICD-0-3/WHO 2008=thyroid; ICD-03 histologies comprising 8050、 8330-8339、 and 8340-8344; and the year of diagnosis from 1975-2016. Only patients > 60 years of age were considered. The study was granted exempt status by our Institutional Review Board. Moreover, the following patients were excluded: death prior to the recommendation for surgery; diagnosed with DTC at the time of autopsy or on the death certificate only were recommended to undergo surgery, but it was unknown if surgery was performed; had incomplete follow-up evaluations; and were not recommended to undergo surgery. Patients in whom the behavior code, ICD-O-2, was benign were also not included. The enrolled patients were divided into two groups (surgery performed and surgery recommended, but not performed).

**Figure 1.**
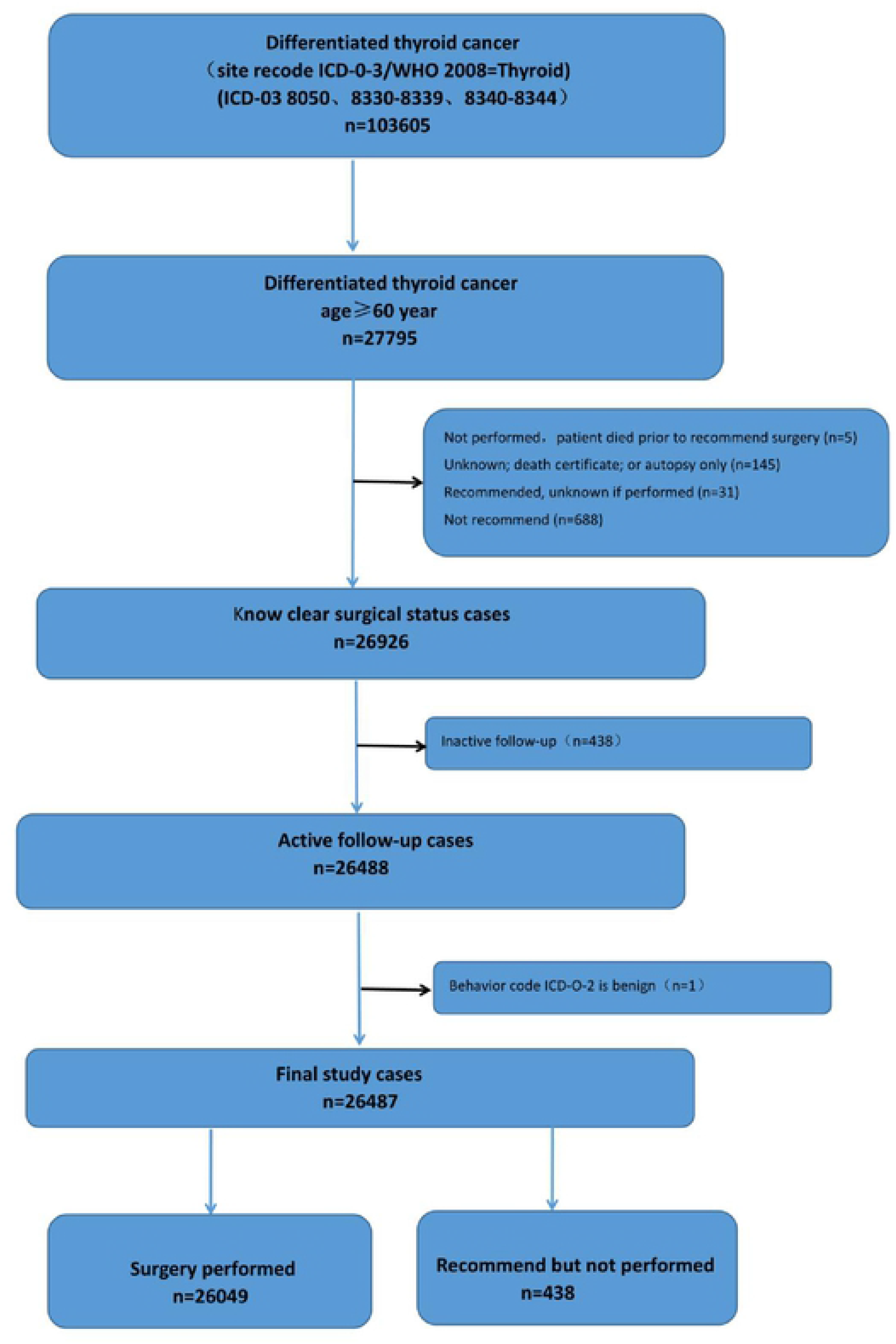
Exclusion criteria utilized to derive the final study cohort from the SEER database.

Furthermore, we also aimed to identify whether post-operative RAI-131 therapy had an impact on CSS with different TNM stages and the number of positive lymph nodes. Thus, we divided patients with T4, N1b, M1, T1-3N0-1a, and the number of metastatic lymph nodes who underwent a total or near-total thyroidectomy.

### Outcome Measures

The outcomes of interest included thyroid CSS and duration of survival for males and females. Duration of survival was defined as the time from the date of diagnosis to the date of death or last contact. Cause of death was defined using the SEER cause of death recode. Patients who died not due tothyroid carcinoma were designated as CSS while patients diagnosed with thyroid carcinoma who were still alive as overall survival.

### Statistical analyses

Mean, and interquartile ranges were reported for continuous variables. Frequency and proportion were reported for categorical variables. Pearson *χ*^2^ test or Fisher’s exact test were used to compare means and proportions, respectively, between categorical variables and treatment groups. The Kaplan Meier estimator was used to determine thyroid cancer CSS and plot survival curves. Cox proportional hazards regression was used to determine hazard ratios (HRs) with 95% confidence intervals (CIs). All statistical tests were two-sided and statistical significance was defined as a P < 0.05.

## RESULTS

### Patient characteristics

Overall, 26,487 elderly patients (≥ 60 years of age) diagnosed with DTC were identified; 26,049 patients underwent surgical treatment and 438 were recommended to undergo surgery, but surgery was not performed. The mean age at the time of diagnosis was 73.50 (SD ± 7.07) years, 68.97 (SD ± 6.90) years in the group that had surgery, and 76 (SD ± 8.64) years in the group that was recommended to have surgery, but did not undergo surgery (Table 1).

**Table 1.** Baseline demographic and clinical features of total patients in this study.

| Characteristics | Surgery Performed<br>( n=26049 ) | Recommended but not<br>surgery(n=438) |
| --- | --- | --- |
| Age at diagnosis<br>(mean±SD) | 68.98±6.90 | 73.56±8.64 |
| 60-64year | 8273(31.76%) | 82(18.72%) |
| 65-69year | 7211(27.68%) | 72(16.44%) |
| 70-74year | 4979(19.11%) | 91(20.71%) |
| 75-79year | 3300(12.67%) | 70(15.98%) |
| 80-85year | 1606(6.17%) | 65(14.84%) |
| 85+year | 680(2.61%) | 58(13.24%) |
| Sex |  |  |
| Male | 7987(30.66%) | 149(34.02%) |
| Female | 18062(69.37%) | 289(65.98%) |
| Race |  |  |
| Black | 1987(7.63%) | 29(6.62%) |
| White | 21606(82.94%) | 349(79.68%) |
| Other | 2316(8.89%) | 53(12.10%) |
| Unknown | 140(0.53%) | 7(1.60%) |
| Surgery Extent |  |  |
| Surgery | 26049(100%) | 0(0.00%) |
| No Surgery | 0(0.00%) | 438(100%) |

In the group that had surgery, 463 and 200 T4/N1b/M1 (7^th^ American Joint Committee on Cancer [AJCC]) patients did and did not receive RAI-131 therapy, respectively. Of 2548 patients, 223 (not T1-3N0) underwent total or subtotal thyroidectomy and 209 and 132 T1-3N1a patients underwent total or subtotal thyroidectomy, respectively. In these cohorts, the mean ages were 69.52 (SD ± 7.18) versus 71.87 (SD± 8.25) years, 67.90 (SD ± 6.11) versus 67.91 (SD± 6.36) years, and 67.88 (SD ± 6.64) versus 67.77 (SD± 6.54) years between patients who did or did not receive RAI-131 therapy, respectively (Table 2).

**Table 2.** different TNM stage patients of unerogone RAI-131 therapy

| Characteristics<br>(AJCC 7th) | RAI-131 therapy<br>(n=3220) | Not RAI-131 therapy<br>(n=555) |
| --- | --- | --- |
| Age of T4/N1b/M1<br>(mean ± SD) | 69.52 ± 7.18 | 71.87 ± 8.25 |
| Male | 215(6.68%) | 82(14.77%) |
| Female | 248(7.70%) | 118(21.26%) |
| Age of T1-3N0<br>(mean ± SD) | 67.90 ± 6.11 | 67.91 ± 6.36 |
| Male | 793(24.63%) | 50(9.01%) |
| Female | 1755(54.50%) | 173(31.17%) |
| Age of T1-3N1a<br>(mean ± SD) | 67.88 ± 6.64 | 67.77 ± 6.54 |
| Male | 76(2.36%) | 59(10.63%) |
| Female | 133(4.13) | 73(13.15%) |

### Survival outcomes

The 120-month CSS rate of females (96.27%-81.92%) and males (95.54%-79.86%) showed a gradual declining trend from patients 60-64 to ≥80 years of age and stage in the group that had surgery. The CSS rate of females (90.68%-52.00%) and males (94.74%-16.22%) showed a marked downward and irregular trend with an increased age in the group recommended to have surgery, but who did not have surgery.

Univariate analysis indicated that the surgery group had a higher 120-month CSS among females in most stages (60-64 years HR: 0.28, 95% CI: 0.06-7.74, P<0.0001; 65-69 years (HR: 0.46, 95% CI: 0.09-2.44, P=0.1755; 70-74 years HR: 0.36, 95% CI: 0.12-1.06, P=0.0017; 75-79 years HR: 0.29, 95% CI: 0.08-1.04, P=0.0003; ≥80 years HR: 0.26, 95% CI: 0.12-0.57, P<0.0001; Figure 2) and males (60-64 years HR: 0.64, 95% CI: 0.06-7.44, P=0.1958; 65-69 years HR: 0.29, 95% CI: 0.05-1.77, P=0.0003; 70-74 years HR: 0.24, 95% CI: 0.03-1.73, P=0.0022; 75-79 years HR: 0.08, 95% CI: 0.01-0.58, P<0.0001; ≥80 years HR: 0.21, 95% CI: 0.06-0.68, P<0.0001), compared with the group that did not have surgery (Figure 3).

**Figure 2.**
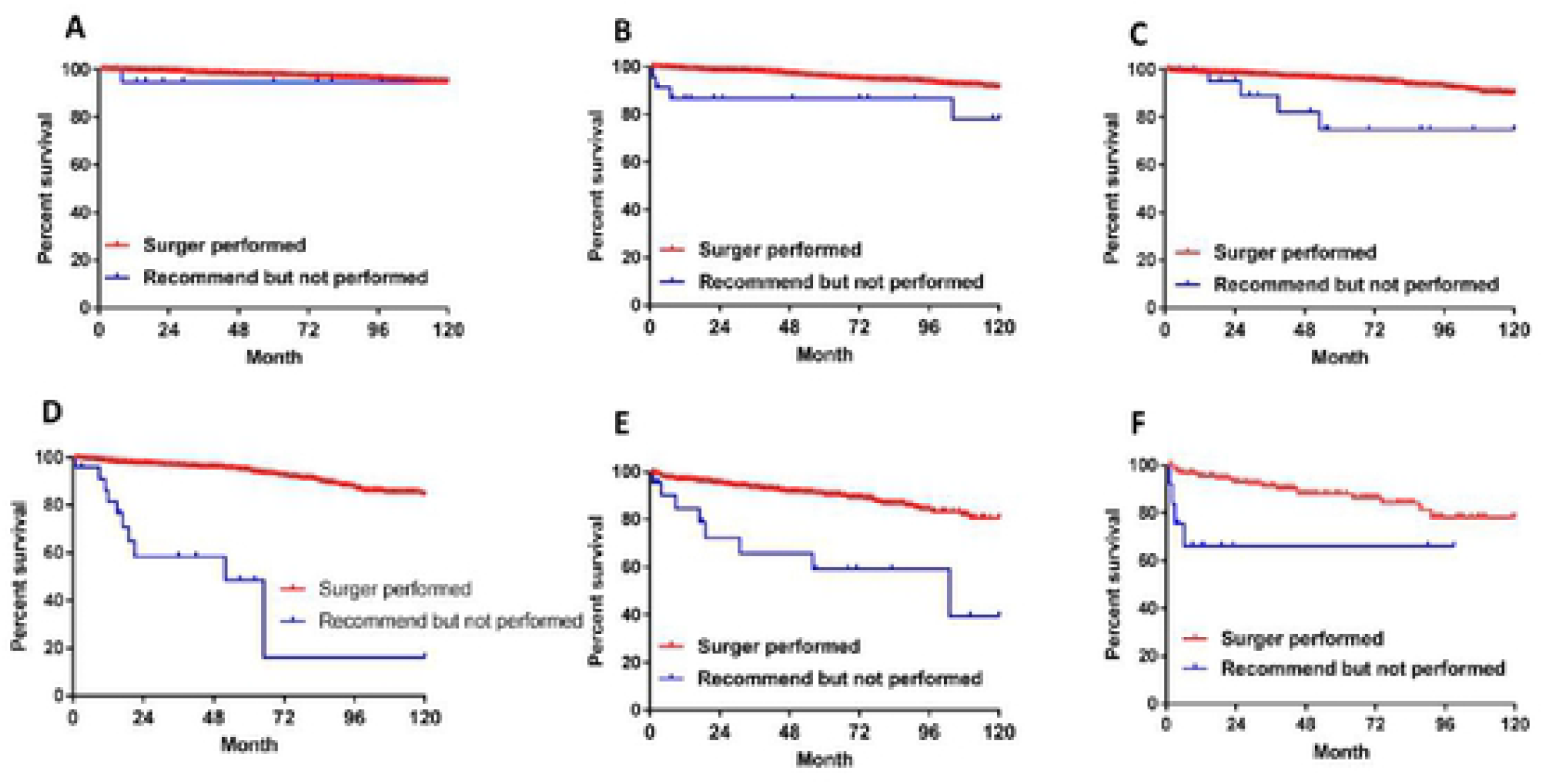
Male Kaplan-Meier survival curve of cause-specific survival comparing the surgery group with the group recommended to have surgery, but did not have surgery of different ages and stages. (A). 60-64 years of age. (B) 65-69 years of age. (C). 70-74 years of age. (D). 75-79 years of age. (E). 80-84 years of age. (F). ≥85 years of age.

**Figure 3.**
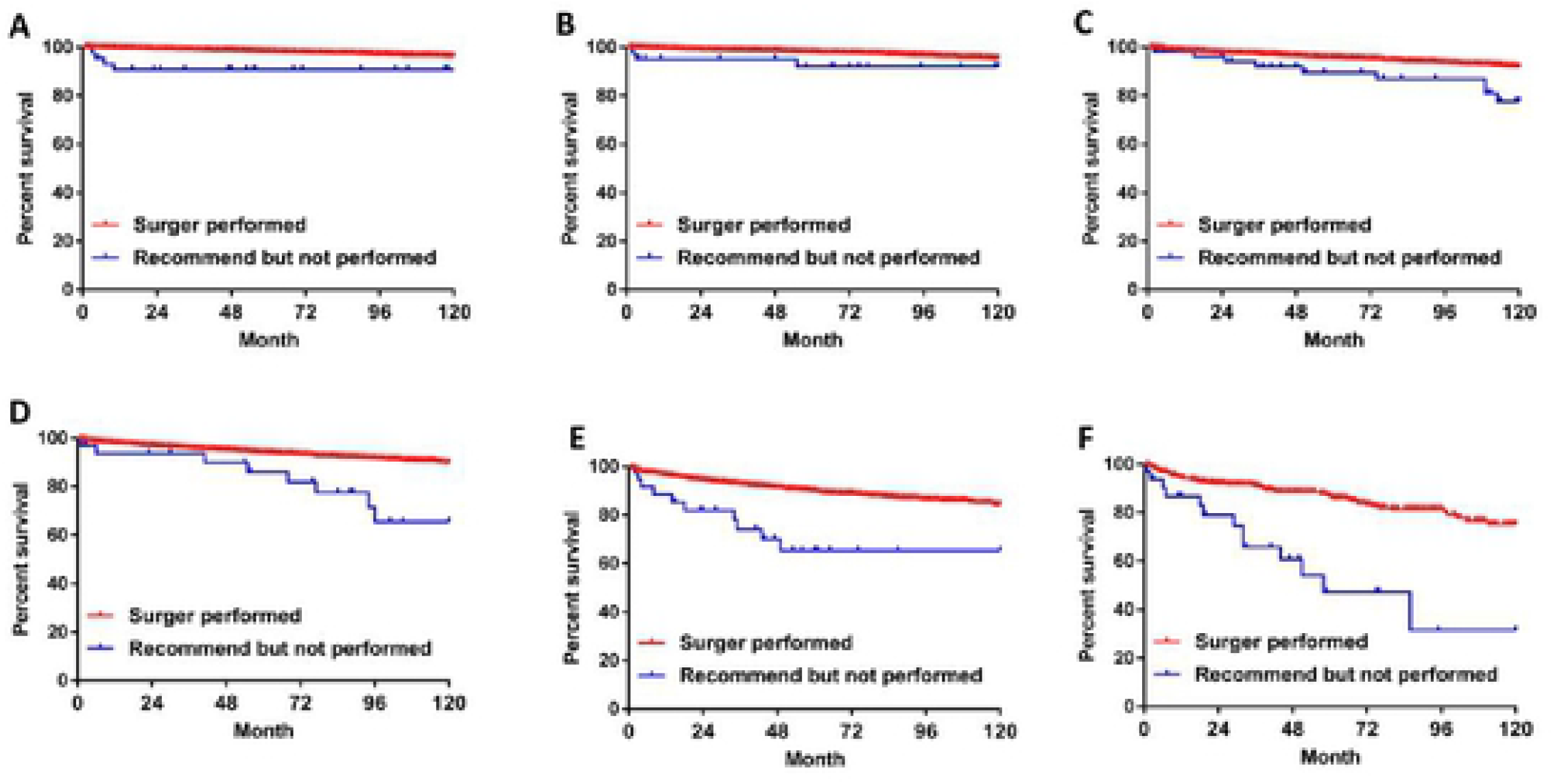
Female Kaplan-Meier survival curve of cause-specific survival comparing the surgery group with the group recommended to have surgery, but did not have surgery of different ages and stages. (A). 60-64 years of age. (B) 65-69 years of age. (C). 70-74 years of age. (D). 75-79 years of age. (E). 80-84 years of age. (F). ≥85 years of age.

RAI-131 therapy was associated with an improved 80-month CSS in T4/N1b/M1 (AJCC 7th) females (HR: 0.47, 95% CI: 0.24-0.95, P<0.0183) and males (HR: 0.27, 95% CI: 0.09-0.84, P<0.0011; Figure 4). There was no differences in CSS among T1-3N0 (P=0.2976, P=0.1650) and T1-3N1a female and male patients (P=0.1040, P=0.5954; Figure 5A-D). Furthermore, we also determined whether the specific number of positive lymph nodes (PLNs) in T1-3N1a patients had different effects on the 80-month CSS, so we divided these patients into subgroups with 1-3 and >3 PLNs; there was no statistical significance between the two subgroups (P=0.0614, P >0.9999; Figure 5E-F).

**Figure 4.**
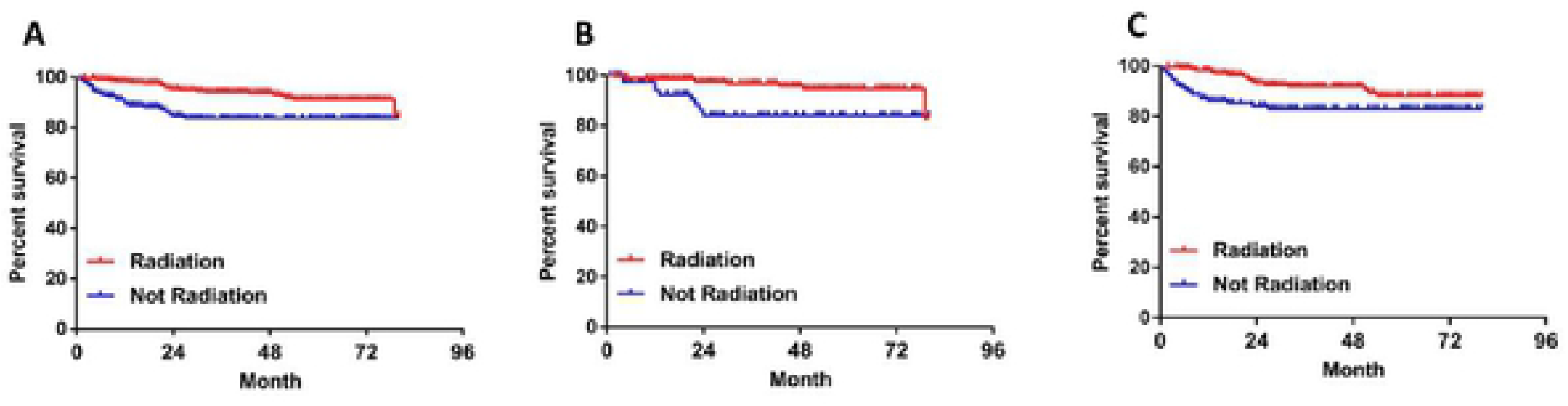
Kaplan-Meier survival curve of cause-specific survival comparing patients who received RAI-131 therapy with high-risk patients who did not receive RAI-131 therapy. (A). T4/N1b/M1 patients (7^th^ AJCC). (B). Male T4/N1b/M1 patients (7^th^ AJCC). (C). Female T4/N1b/M1 patients (7^th^ AJCC).

**Figure 5.**
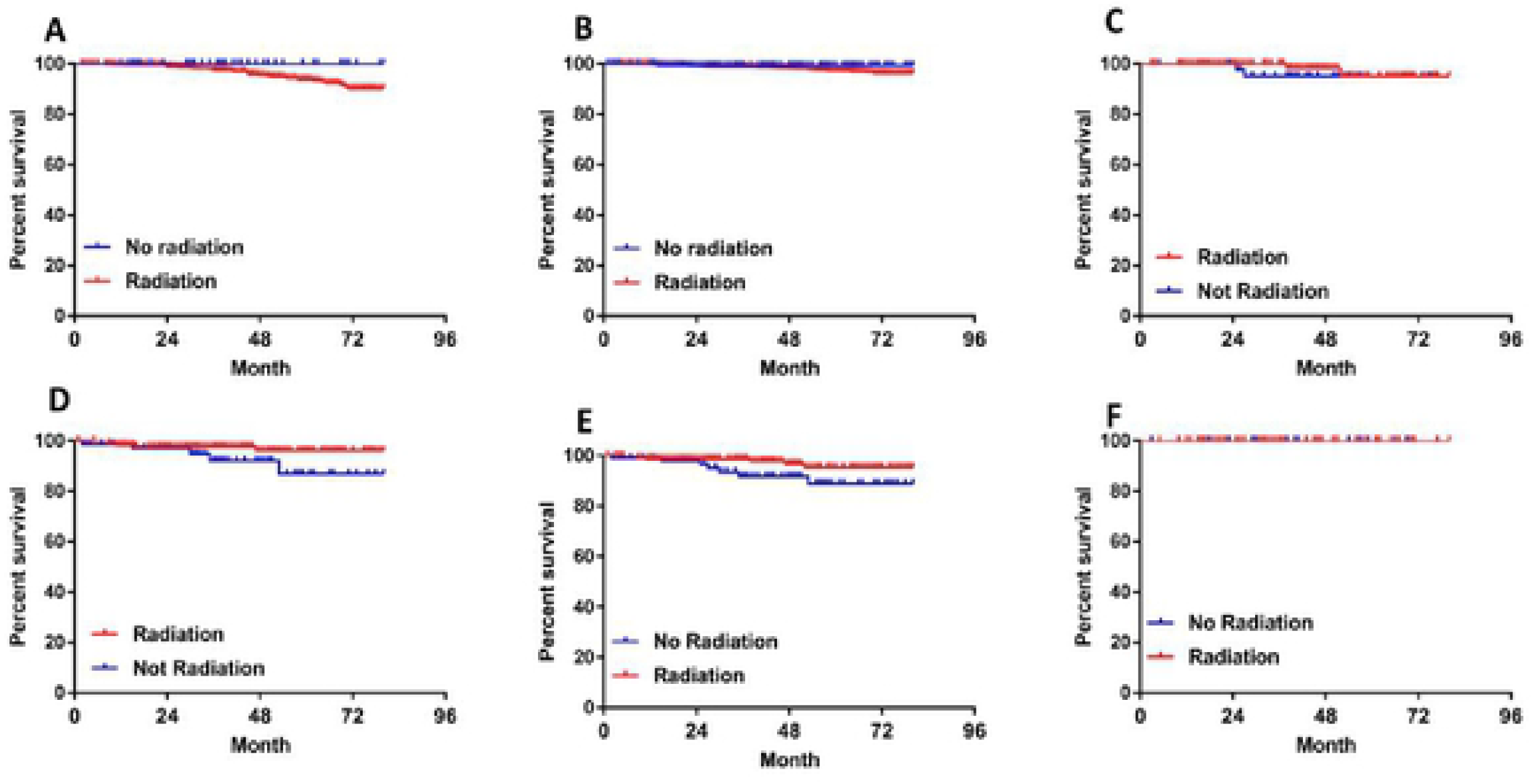
Kaplan-Meier survival curve of cause-specific survival comparing T1-3N0-1a (7^th^ AJCC) patients who did and did not receive RAI-131 therapy. (A). T1-3N0 male patients. (B). T1-3N0 female patients. (C). T1-3N1a male patients. (D). T1-3N1a female patients. (E). T1-3N1a patients with 1-3 lymph node metastases. (F). T1-3N1a patients with > 3 lymph node metastases.

## DISCUSSION

DTC is the only human malignancy to include age as a part of the AJCC staging system, with a distinct cut point at 55 years of age [8]. Older age was also associated with a greater probability of a death from other reasons. Therefore, the benefit and risk of surgery should be considered as part of a balanced clinical approach due to the possibility of death from both DTC and other causes in older patients. It is empirically clear that many older patients, especially patients with smaller thyroid cancers, do not progress and if carefully triaged are otherwise eligible for monitoring. The individual acceptance of risk (fro m cancer or from surgery) thus remains a matter of informed consensus b etween patients and clinician [9,10].

Age has been identified as a well-recognized prognostic determinant of CSS in older patients, although DTC is relatively indolent. Age has a linear dose-dependent relationship with the DTC mortality rate. An age-specific cutoff corresponding to a marked decrement in survival is not apparent. Therefore, DTC patients have a higher cancer-specific mortality rate in elderly patients, rather than the tumor becoming more inactive as age increases [11,12]. Another reason DTC-related mortality accounts for only a fraction of the overall deaths among patients who underwent surgery is that co-morbidities increased overall mortality by increasing the probability of other causes of mortality. Indeed patients died earlier from other co-morbidities and had a low probability of mortality from DTC. The roles of co-morbidities and mortality from other causes were more pronounced in patients with a lower stage of DTC [13–15]. To eliminate these distractions, we only explored the CSS of patients from the SEER database. In the current study, we concluded that older DTC patients, especially men and women > 70 years who underwent thyroidectomy, had a better 10-year CSS. Such patients acquired a longer survival time through surgery if there were no additional high risk diseases associated with death.

In fact, some patients were and were not recommended to undergo surgery because of other diseases or complications.. In the current study only the patients recommended to undergo surgery, but who declined surgery were identified as the no surgery group and the effect of other diseases and complications were excluded. Therefore, the results were more favorable for the treatment strategy of elderly patients with DTC.

The 2015 ATA guidelines suggest modifications in risk stratification for DTC patients. The consensus guidelines recommend that RAI-131 should be offered to high-risk patients to reduce the risk of disease recurrence or mortality for DTC patients after a total or near-total thyroidectomy [16]; however, inappropriate use of RAI places patients at unnecessary risk of permanent treatment-related toxicity and secondary cancers[17]. Post-operative adjuvant RAI therapy has become more selective based on current guidelines [4,18].

The updated guidelines have supported both decreased RAI doses for select populations, as well as expanded definitions for low- and intermediate-risk patients that may not require RAI-131. A growing body of literature indicates that the benefit-to-risk ratio is necessary to accurately identify patients who are ideal candidates for the therapy. RAI-131 can avoid unnecessary treatment, cost, and adverse effects at the same time [4,16,19,20]. Evidence surrounding the use of RAI-131 in patients with intermediate-risk DTC is less robust and there is no consensus regarding the benefit-to-risk ratio [21].

The debate centers on the question of appropriate use of post-operative adjuvant RAI therapy. T4/N1b/M1 (7^th^ AJCC) patients who received RAI-131 therapy had a better CSS than patients who did not RAI-131 therapy; however, there was no difference in the number of PLNs (0, 1-3, and >3) among T1-3N0-1a patients (7^th^ AJCC), which are stage I or II (8^th^ AJCC). There was no difference in T1-3N0-1a patients with 0, 1-3, and >3 PLNs. Therefore, the patients at high-risk have a higher benefit-to-risk ratio by RAI-131 therapy. Otherwise, the intermediate- or low-risk patients have fewer benefits from RAI therapy, thus, they achieve equal survival with thyroid stimulating hormone-suppressive therapy and active follow-up without RAI-131 therapy and also could avoid the associated risks of adverse effects.

The endocrinologist and nuclear medicine physician have an indispensable role in RAI-131 decision-making, the role of surgeons has been shown to exert substantial indirect influence on RAI-131 use, and the treatment concept is often a pivotal factor in post-operative management, especially for intermediate- and low-risk patients. Such patterns, however, may lead to inappropriate use of RAI-131. The ultimate goal of this study was to selectively leverage the strengths of RAI-131 therapy and determine whether use of RAI-131 is suitable for high-risk patients who benefit, as well as intermediate- and lower-risk patients who may not benefit based on a multidisciplinary, risk-adapted approach [22].

There were several limitations to the current study. This was a non-randomized observational study with the possibility of selection bias and inherent coding errors, especially an inability to confirm histology status in patients who did not undergo surgery. The SEER database did not capture key information variables, such as the largest size of metastatic lymph nodes, extranodal extension, vascular invasion, the histologic subtype, genetic mutations, specific RAI-131 dose, and other tumor or nodal factors that may have influenced on the decision to administer RAI-131 treatment, cancer recurrence, and prognosis outcomes. The number of patients recommended to undergo surgery, but who did not have surgery was small, and the TNM status of most patients was blank, so we cannot further analyze the influence of different TNM staging on CSS.

In conclusion, our analysis indicated that surgical treatment should be recommended to elderly DTC patients because surgery can help achieve a better CSS. The high-risk patients have a greater benefit-to-risk ratio through RAI-131 therapy. To avoid the related adverse effects associated with RAI-131 therapy, a multidisciplinary discussion should take place for intermediate- and low-risk patients.

## References

1. Davies L, Welch HG. Current thyroid cancer trends in the United States. Jama Otolaryngol Head Neck Surg. 2014;140(4):317–322. https://doi:10.1001/jamaoto.2014.1 PMID:24557566

2. Fahiminiya S, de Kock L, Foulkes WD. Biologic and Clinical Perspectives on Thyroid Cancer. N Engl J Med. 2016;375(23):2306–2307. https://doi:10.1056/NEJMc1613118 PMID:27959678

3. Aschebrook-Kilfoy B, Kaplan EL, Chiu BC, Angelos P, Grogan RH. The acceleration in papillary thyroid cancer incidence rates is similar among racial and ethnic groups in the United States. Ann Surg Oncol. 2013;20(8):2746–2753. https://doi:10.1245/s10434-013-2892-y PMID:23504142

4. Haugen BR, Alexander EK, Bible KC, et al. 2015 American Thyroid Association Management Guidelines for Adult Patients with Thyroid Nodules and Differentiated Thyroid Cancer: The American Thyroid Association Guidelines Task Force on Thyroid Nodules and Differentiated Thyroid Cancer. Thyroid. 2016;26(1):1–133. https://doi:10.1089/thy.2015.0020 PMID:26462967

5. Jeong SY, Lee SW, Kim WW, Jung JH, Lee WK, et al. Clinical outcomes of patients with T4 or N1b well-differentiated thyroid cancer after different strategies of adjuvant radioiodine therapy. Sci Rep. 2019;9(1): 5570. https://doi:10.1038/s41598-019-42083-3 PMID:30944403

6. National Cancer Institute. Surveillance Epidemiology and End Results (SEER) Program, Available at:http://seer.cancer.gov/popdata/methods.html. Accessed: January 10, 2017; National Cancer Institute: Surveillance, Epidemiology,and End Results. http://www.seer.cancer.gov.

7. Surveillance, Epidemiology, and End Results Program (SEER). Available at: www.seer.cancer.gov. Accessed 29 January 2018.

8. Amin MB, Greene FL, Edge SB, Compton CC, Gershenwald JE, et al. The Eighth Edition AJCC Cancer Staging Manual: Continuing to build a bridge from a population-based to a more “personalized” approach to cancer staging. CA Cancer J Clin. 2017;67(2):93–99. https://doi:10.3322/caac.21388 PMID:28094848

9. Ho AS, Luu M, Zalt C, Morris LGT, Chen I, et al. Mortality Risk of Nonoperative Papillary Thyroid Carcinoma: A Corollary for Active Surveillance. Thyroid. 2019;29(10):1409–1417. https://doi:10.1089/thy.2019.0060 PMID:31407637

10. Miyauchi A, Kudo T, Ito Y, Oda H, Sasai H, et al. Estimation of the lifetime probability of disease progression of papillary microcarcinoma of the thyroid during active surveillance. Surgery. 2018;163(1) 48–52. https://doi:10.1016/j.surg.2017.03.028 PMID:29103582

11. Adam MA, Thomas S, Hyslop T, Scheri RP, Roman SA, et al. Exploring the Relationship Between Patient Age and Cancer-Specific Survival in Papillary Thyroid Cancer: Rethinking Current Staging Systems. J Clin Oncol. 2016;34(36):4415–4420. PMID:27998233

12. Ganly I, Nixon IJ, Wang LY, Palmer FL, Migliacci JC, et al. Survival from Differentiated Thyroid Cancer: What Has Age Got to Do with It? Thyroid. 2015;25(10):1106–1114. https://doi:10.1089/thy.2015.0104 PMID: 26148759

13. Lee YK, Hong N, Park SH, Shin DY, Lee CR, et al. The relationship of comorbidities to mortality and cause of death in patients with differentiated thyroid carcinoma. Sci Rep. 2019;9(1):11435. https://doi:10.1038/s41598-019-47898-8 PMID:31391492

14. Leite AK, Kulcsar MA, de Godoi Cavalheiro B, de Mello ES, Alves VA, et al. DEATH RELATED TO PULMONARY METASTASIS IN PATIENTS WITH DIFFERENTIATED THYROID CANCER. Endocr Pract. 2017;23(1):72–78. https://doi:10.4158/EP161431 PMID:27749128

15. Gillanders SL, O’Neill JP. Prognostic markers in well differentiated papillary and follicular thyroid cancer (WDTC). Eur J Surg Oncol. 2018;44(3):286–296. https://doi:10.1016/j.ejso.2017.07.013 PMID:28801060

16. Sun Y, Gong J, Guo B, Shang J, Cheng Y, et al. Association of adjuvant radioactive iodine therapy with survival in node-positive papillary thyroid cancer. Oral Oncol. 2018;87:152–157. https://doi:10.1016/j.oraloncology.2018.10.041 PMID:30527231

17. Marti JL, Davies L, Haymart MR, Roman BR, Tuttle RM, et al. Inappropriate Use of Radioactive Iodine for Low-Risk Papillary Thyroid Cancer Is Most Common in Regions with Poor Access to Healthcare. Thyroid. 2015;25(7):865–866. https://doi:10.1089/thy.2015.0112 PMID:25963001

18. Luster M, Aktolun C, Amendoeira I, Barczyński M, Bible KC, et al. European Perspective on 2015 American Thyroid Association Management Guidelines for Adult Patients with Thyroid Nodules and Differentiated Thyroid Cancer: Proceedings of an Interactive International Symposium. Thyroid. 2019;29(1):7–26. https://doi:10.1089/thy.2017.0129 PMID:30484394

19. Adam MA, Pura J, Goffredo P, Dinan MA, Reed SD, et al. Presence and Number of Lymph Node Metastases Are Associated With Compromised Survival for Patients Younger Than Age 45 Years With Papillary Thyroid Cancer. J Clin Oncol. 2015;33(21):2370–2375. https://doi:10.1200/JCO.2014.59.8391 PMID:26077238

20. Asimakopoulos P, Nixon IJ, Shaha AR. Differentiated and Medullary Thyroid Cancer: Surgical Management of Cervical Lymph Nodes. Clin Oncol (R Coll Radiol). 2017;29(5):283–289. https://doi:10.1016/j.clon.2017.01.001 PMID:28094086

21. Orosco R, Hussain T, Noel JE, Chang DC, Dosiou C, et al. Radioactive iodine in differentiated thyroid cancer: a national database perspective. Endocr Relat Cancer. 2019. https://doi:10.1530/ERC-19-0292 PMID:31443087

22. Marti JL, Morris LGT, Ho AS. Selective use of radioactive iodine (RAI) in thyroid cancer: No longer “one size fits all”. Eur J Surg Oncol. 2018;44(3):348–356. https://doi:10.1016/j.ejso.2017.04.002 PMID:28545679

